# Differential covariance: A new method to estimate functional connectivity in fMRI

**DOI:** 10.1101/2020.06.16.155630

**Authors:** Tiger W. Lin, Yusi Chen, Qasim Bukhari, Giri P. Krishnan, Maxim Bazhenov, Terrence J. Sejnowski

**Affiliations:** Neurosciences Graduate Program, University of California San Diego, La Jolla, CA, 92092; Computational Neurobiology Laboratory, Salk Institute for Biological Sciences, La Jolla, CA, 92037; Institute for Neural Computation, University of California San Diego, La Jolla, CA, 92092; Division of Biological Sciences, University of California San Diego, La Jolla, CA, 92092; McGovern Institute for Brain Research, Massachusetts Institute of Technology, Cambridge, MA, 02139; Department of Medicine, University of California San Diego, La Jolla, CA, 92092

**Keywords:** Differential covariance, Functional connectivity, fMRI, Network modeling

## Abstract

Measuring functional connectivity from fMRI recordings is important in understanding processing in cortical networks. However, because the brain’s connection pattern is complex, currently used methods are prone to producing false functional connections. We introduce differential covariance analysis, a new method that uses derivatives of the signal for estimating functional connectivity. We generated neural activities from Dynamical Causal Modeling and a neural network of Hodgkin-Huxley neurons and then converted them to hemodynamic signals using the forward Balloon model. The simulated fMRI signals together with the ground truth connectivity pattern were used to benchmark our method with other commonly used methods. Differential covariance achieved better results in complex network simulations. This new method opens an alternative way to estimate functional connectivity.

## 1 Introduction

*Analysis of fMRI recordings to extract functional connectivity aims to characterize a pattern of statistical associations among time series measured from different brain regions (Reid et al., 2019). It has been used quite successfully in identifying brain network interactions at rest and during certain behaviors (Raichle, 2015).* Moreover, interesting geometric properties of brain networks can be studied with this method (Achard et al., 2006).

*Many statistical methods have been proposed for estimating functional connectivity, including covariance-based methods, lag-based methods (Seth, 2010; Geweke, 1984) and Bayes-net methods (Mumford and Ramsey, 2014). However, without knowing the ground truth, it has been difficult to assess the validity of these methods.* To compare these methods, we simulated BOLD signals from network models with known patterns of connectivity (Friston et al., 2003). The simulated neural signals were then transformed to BOLD sgnals using an fMRI forward model using the nonlinear balloon model (Buxton et al., 1998). In a previous study (Smith et al., 2011), many well known statistical methods were benchmarked with simulated fMRI datasets, and concluded that covariance-based methods performed best.

As previously reviewed in Stevenson et al. (2008), a fundamental problem for covariance-based methods is that two correlated nodes do not necessarily have a direct physical connection. *This could be problematic when one wants to compare the functional connectivity with the anatomical wiring of the brain such as fiber tracts or neural projections.*

We introduce a new method to reduce these false connections and provide estimates that are closer to ground truth in simulated fMRI data. *Our new method is a novel form of functional connectivity with weighted strength that extracts information about the directionality of causal influence. This method, called differential covariance, performs better than other covariance-based methods on synthetic data by taking advantage of differential signals.*

This paper is organized as follows: in Section 2 we explain how synthetic BOLD signals were generated and introduce the new method. We also explain how results from different estimators were quantified; in Section 3, we compare the performance of our new method with the performance of other methods; in Section 4, we discuss the advantages and generalizability of the differential covariance.

## 2 Methods

### 2.1 BOLD signal simulations

#### 2.1.1 Dynamic causal modeling (DCM)

**Figure 1:**
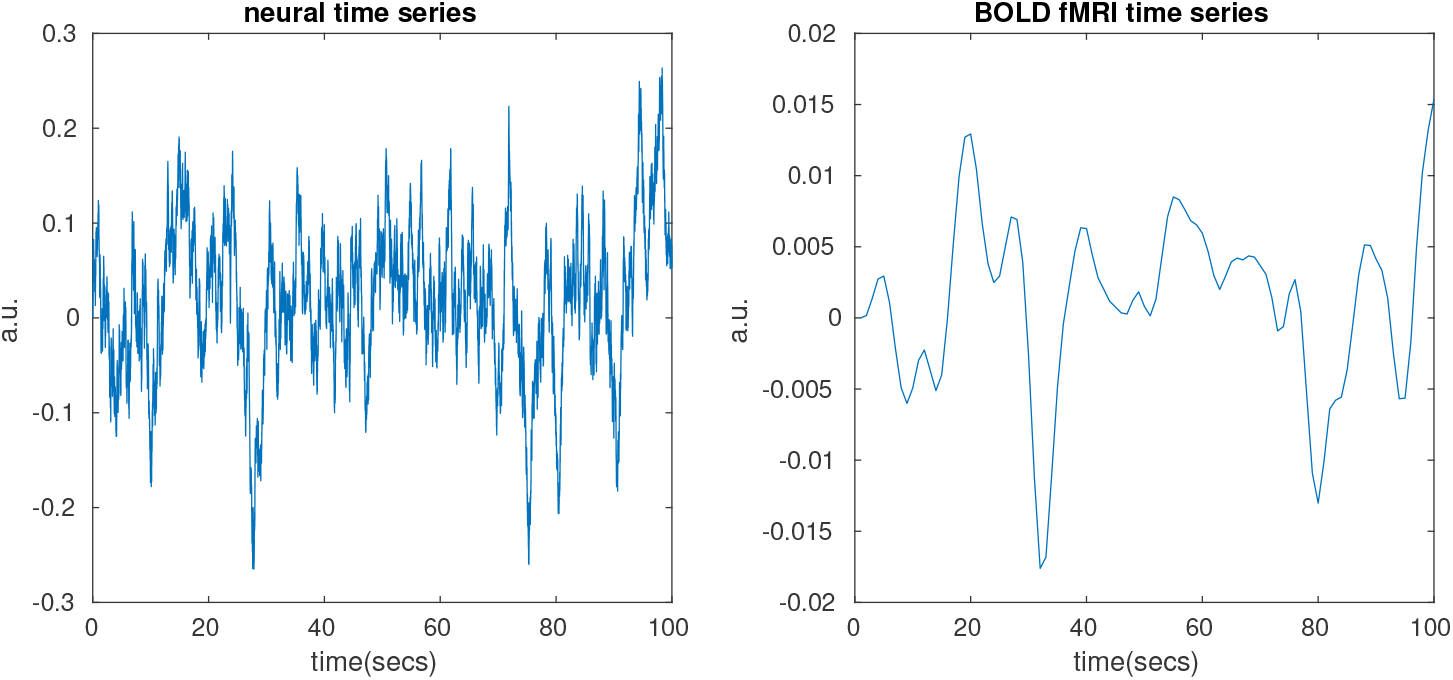
Neural signal and BOLD signal example

We used dynamic causal modeling (DCM) (Friston et al., 2003) and a more realistic thalamocortical model (Details explained in Section 2.1.3) to simulate neural signals. Neural signals were then converted to BOLD signals using the classical forward Balloon model with revised coefficients (Obata et al., 2004). The detailed method of DCM was previously explained in Smith et al. (2011). Briefly, the neural signals were generated from the model:

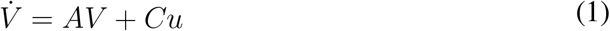

where, *V* are time series of neural signals and *A* is the connection matrix. Following the parameters used in the previous work (Smith et al., 2011), the diagonal values of *A* are assigned to be −1 in our simulations. Its non-diagonal values represent the connection pattern between nodes, which is the functional connectivity we try to estimate. The connection strength is uniformly distributed between 0.4 and 0.6. Figure. 3A shows an example connection pattern represented by a 60 *×* 60 matrix. Its first 50 nodes are visible, and they are connected with a sparse pattern. The 10 additional nodes are latent and have a broad connection to the visible nodes. We added the latent inputs to the system because unobserved common inputs exist in real-world problems (Stevenson et al., 2008). Variable *u* is the external stimuli. For our simulations, we assume there is no external stimuli to the network. However, a white Gaussian noise with 0.1 variance is added to *u*. *C* is an identity matrix in this paper indicating that each node has its independent external stimuli.

**Figure 3:**
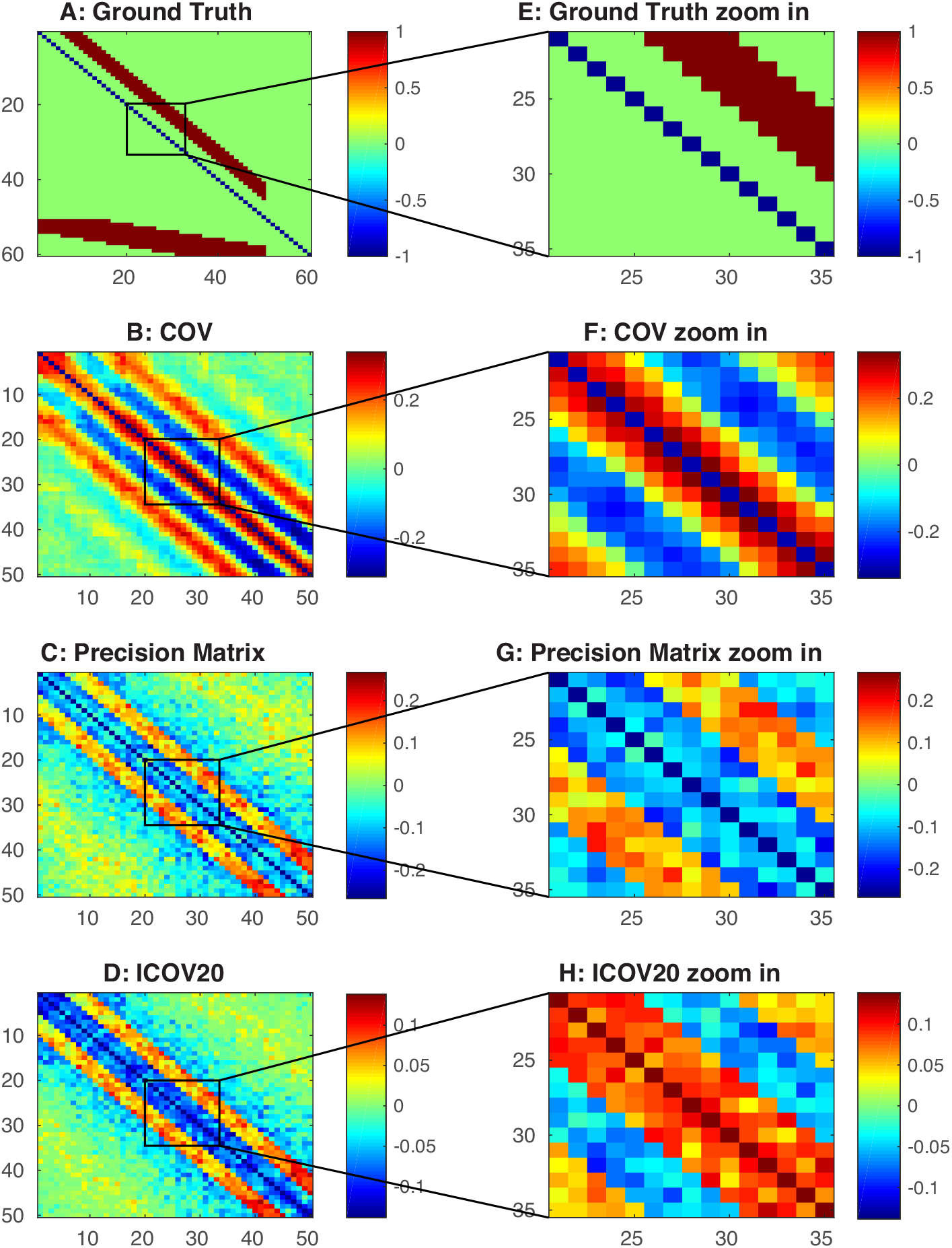
DCM model’s connection ground truth and connectivity estimation from covariance-based methods. A) Ground truth connection matrix. Node No.1-50 are visible nodes. Node No.51-60 are latent nodes. B) Estimation from the covariance method. C) Estimation from the precision matrix method. D) Estimation from the sparsely regularized precision matrix (ICOV) method. E) Zoom in of panel A. F) Zoom in of panel B. G) Zoom in of panel C. H) Zoom in of panel D.

Once the neural signals were simulated from the neural network model, they were fed into the Balloon model to generate BOLD signals. The Balloon model simulates the nonlinear vascular dynamic response due to the neural activities (Obata et al., 2004):

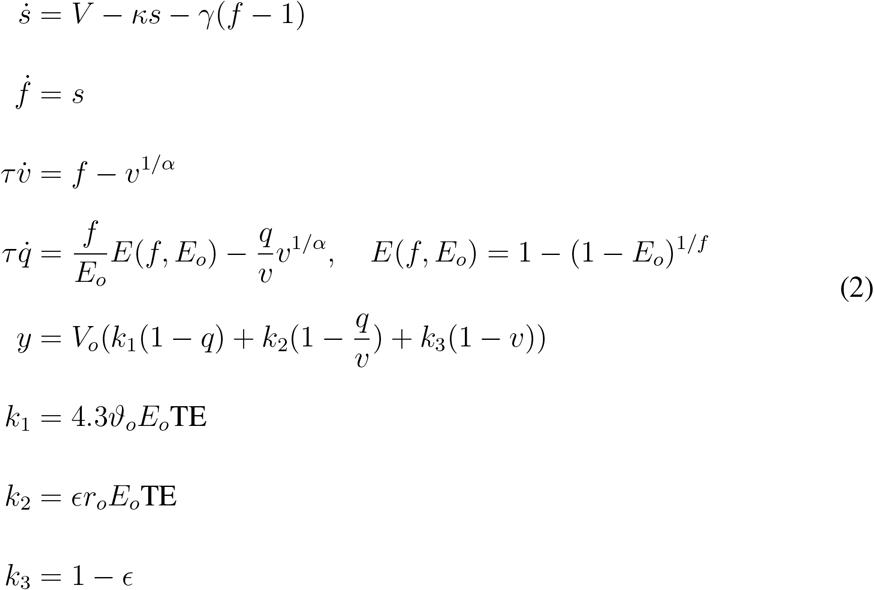

where the superscript dot stands for first order derivative with respect to time, *V* is the neural signal, *s* is the vasodilatory signal, *f* is the blood inflow, *v* is blood volume, *q* is deoxyhemoglobin content and TE is time of echo in seconds. The BOLD signal *y* is a combined signal of the blood volume and the deoxyhemoglobin content. Lastly, the BOLD signal *y* is sampled at 1 sec. Values and notations of the constants and coefficients are provided in Appendix B Table 1.

**Table 1:**
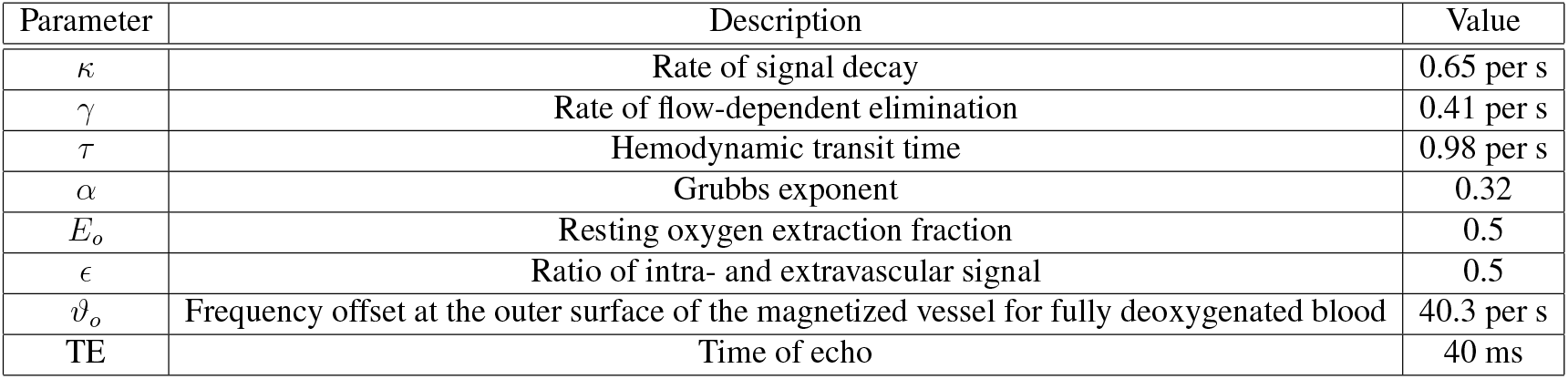
Balloon model parameters

#### 2.1.2 Benchmark dataset from previous work

Based on DCM, using different connection patterns, a benchmark dataset of 28 simulations was previously created (Smith et al., 2011). *We slightly modified the simulation process to evaluate the performance of our method and previous methods over this popular benchmark dataset.*

First, *the original binary external stimulus was replaced by a random Gaussian input because our method cannot deal with sharp state transitions. However, our method is able to give correct estimation in a network with gradual state transition, as in the case of the thalamocortical model simulations (Section 2.1.3).*

Second, we reduced the sampling interval (TR) from 3 seconds to 1 second. This is a reasonable modification because an increasing number of fMRI studies (such as the Human Connectome Project) is adopting a densely sampled protocol with TR less than or equal to 1 second.

Third, because the new method needs more data to provide a good estimation, simulation duration was increased from 600 second to 6000 seconds for most of the simulations.

Fourth, the thermal noise added to the BOLD signal was changed from 1% to 0.1%. All other procedures in the original paper (Smith et al., 2011) were followed. Twentyeight different connection patterns were first simulated using DCM and passed through the forward Balloon model as shown in Equation 2.

#### 2.1.3 Thalamocortical model

To further validate our method, a more realistic Hodgkin-Huxley based ionic neural network model was used in this paper to generate neural signals and the neural signals was then passed through the forward Balloon model (Equation 2) to generate the BOLD signals. The thalamocortical model used in this study was based on several previous studies (Bazhenov et al., 2002; Chen et al., 2012; Bonjean et al., 2011). It was originally used to model spindle and slow wave activity. The thalamocortical model was structured as a one-dimensional, multi-layer array of cells. It consisted of 50 cortical pyramidal (PY) neurons, 10 cortical inhibitory (IN) neurons, 10 thalamic relay (TC) neurons and 10 reticular (RE) neurons. The connection between the 50 PY neurons is shown in Fig. 6A. For the rest of the connection types, a neuron connects to all target neurons within a predefined radius as described in Bazhenov et al. (2002). The details of this model can be found in our previous publication (Lin et al., 2017).

**Figure 6:**
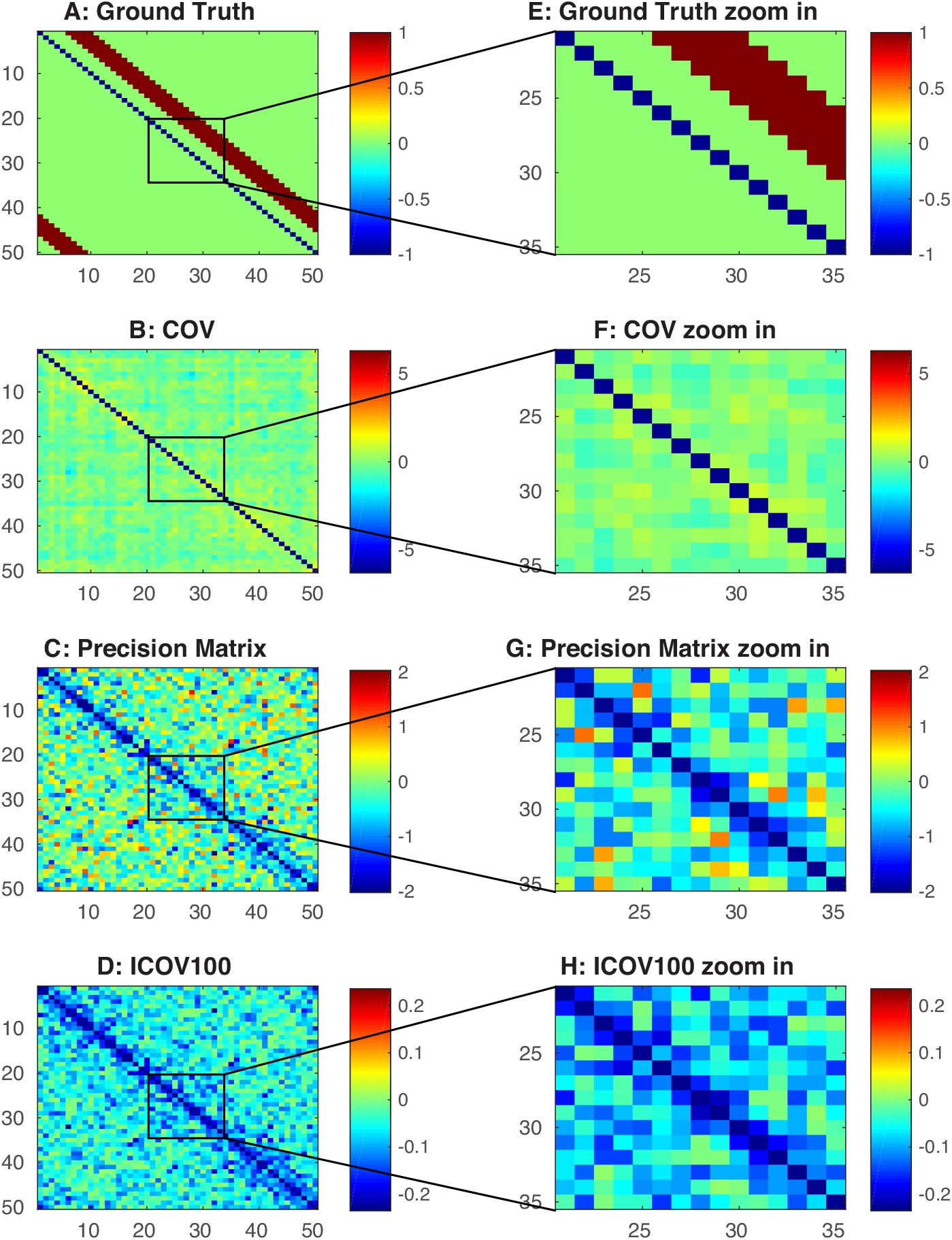
Connectivity estimation from ovariance-based methods of synthetic BOLD signals from thalamocortical model. A) Ground truth connection of the PY neurons in the thalamocortical model. B) Estimation from the covariance method. C) Estimation from the precision matrix method. D) Estimation from the sparsely regularized precision matrix (ICOV) method. E) Zoom in of panel A. F) Zoom in of panel B. G) Zoom in of panel C. H) Zoom in of panel D.

For each simulation, we simulated the network for 4000 seconds with 600 seconds at each sleep stage (Awake→Stage 2→Slow Wave→REM→Stage 2) and 200 seconds of transition period between stages. The simulated neural signals were transferred to BOLD signals and sampled at 10Hz.

### 2.2 New method for functional connectivity estimation

In our new method, we first build a backward model to reconstruct the neural signals from the BOLD signals. Then we applied our differential covariance method to the reconstructed neural signals to estimate the functional connectivity.

#### 2.2.1 Backward model

To derive a backward model, we first linearized the high order terms in the balloon model (Eq. 2) according to Khalidov et al. (2011). To be more specific, let *{x*_1_*, x*_2_*, x*_3_*, x*_4_*}* = *{s,* 1 *− f,* 1 *− v,* 1 *− q}*, and linearize around the resting point *{x*_1_*, x*_2_*, x*_3_*, x*_4_*}* = *{*0, 0, 0, 0*}*, we have:

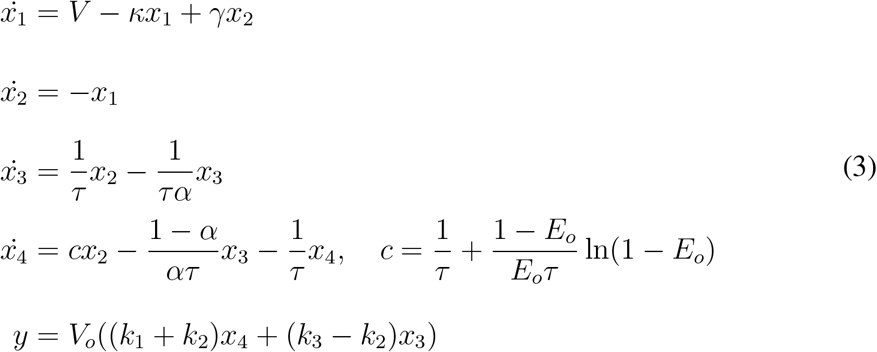

After rearrangement, we have:

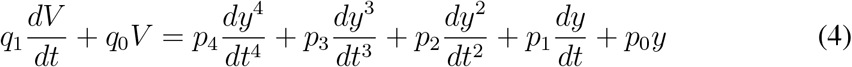

where constants and coefficients are defined in Appendix C. Equation 4 shows that if we compute the high order derivatives of *y*, we can reconstruct the original neuronal signal *V* as a linear combination of its first order derivative and itself. We denote this reconstructed neuronal signal as *z*.

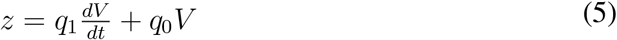

#### 2.2.2 Differential covariance

In our previous work (Lin et al., 2017), we demonstrated that using derivative signals, the differential covariance method can reduce the false functional connections in a simulated network of neuronal activities. Here we briefly explain this method.

First, we assume that using the backward model above, we have reconstructed the neural signals *z*(*t*) of some fMRI nodes. *z*(*t*) is a *N × M* matrix, where *N* is the number of nodes recorded in the network, and *M* is the number of time points during the simulation. We compute the derivative of each time series by calculating 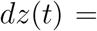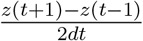. Then, the covariance between *z*(*t*) and *dz*(*t*) is computed and denoted as Δ_*C*_, which is a *N × N* matrix defined as the following:

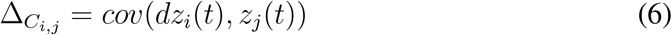

where *z_j_*(*t*) is the reconstructed neural signal of node *j*, *dz_i_*(*t*) is the derivative neural signal of node *i*, and *cov* is the sample covariance function for two time series.

Note that different from our previous work (Lin et al., 2017), the differential covairance is computed using the reconstructed neural signal 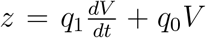 instead of the neural signal *V*, which is not recoverable using the Balloon model.

#### 2.2.3 Partial differential covariance

As previously mentioned in Stevenson et al. (2008), one problem of the covariance method is the propagation of covariance. Here we designed a customized partial covariance algorithm to reduce this type of error in our differential covariance method. We use 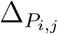 to denote the partial covariance between *dz_i_*(*t*) and *z_j_*(*t*).

Let *Z* be a set of all nodes except *i* and *j*: *Z* = *{*1, 2*, …, i −* 1*, i* + 1*, … j −* 1*, j* + 1*, …, N}*. Using the derivation from Appendix Section A.1.2, we have:

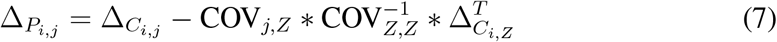

where 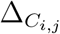 and 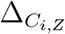 were computed from the previous section, and COV is the covariance matrix of the nodes.

#### 2.2.4 Sparse latent regularization

Finally, we applied the sparse latent regularization method to our estimation (Chandrasekaran et al., 2011; Yatsenko et al., 2015). As explained in Appendix A.4, during recording, there are observable nodes and latent nodes in a network. If the connections between the observable nodes are sparse and the number of latent nodes is small, this method can separate the covariance matrix into these two parts and one can regard the sparse matrix as the intrinsic connections between the observable nodes.

Here we define Δ_*S*_ as the sparse intrinsic connections from the observed nodes, and *L* as the residual covariance part introduced from the latent inputs. Then by solving:

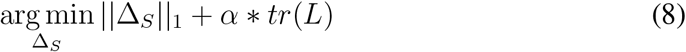

under the constraint that

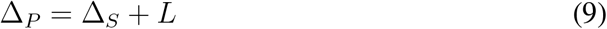

we retrieve the intrinsic connections between the nodes measured. Where, *|| ||*_1_ is the element-wise L1-norm of a matrix, and *tr*() is the trace of a matrix. *α* is the penalty ratio between the L1-norm of Δ_*S*_ and the trace of L. It was set to 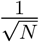 for all our estimations. Δ_*P*_ is the partial differential covariance computed from section 2.2.3.

### 2.3 Performance quantification

The performance of each method is judged by 4 quantified values. The first 3 values indicate the method’s abilities to reduce the 3 types of false connections (Fig. 2). The last one indicates the method’s ability to correctly estimate the true positive connections against all possible interference.

**Figure 2:**
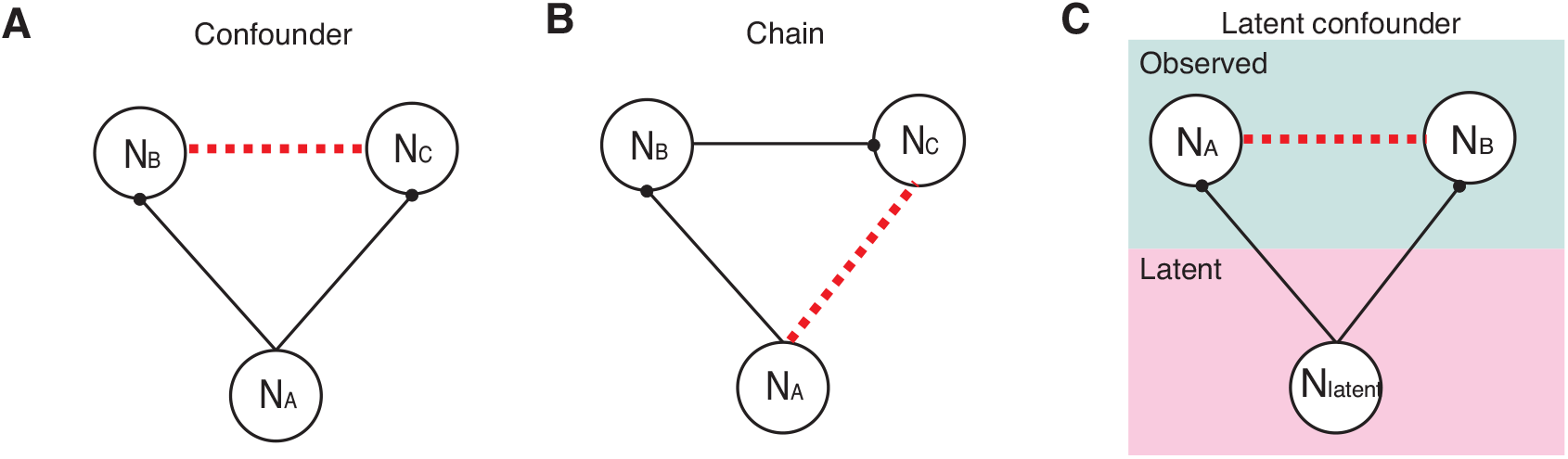
Illustrations of the 3 types of false connections in the covariance-based methods. Solid lines indicate the physical wiring between neurons, and the black circles at the end indicate the synaptic contacts (i.e. the direction of the connections). The dotted lines are the false connections introduced by the covariance-based methods. A) Confounder errors, which are due to two neurons receiving the same synaptic inputs. B) Chain errors, which are due to the propagation of correlation. C) Latent confounder errors, which are due to unobserved common inputs.

Let’s define *G* as the ground truth connectivity matrix, where:

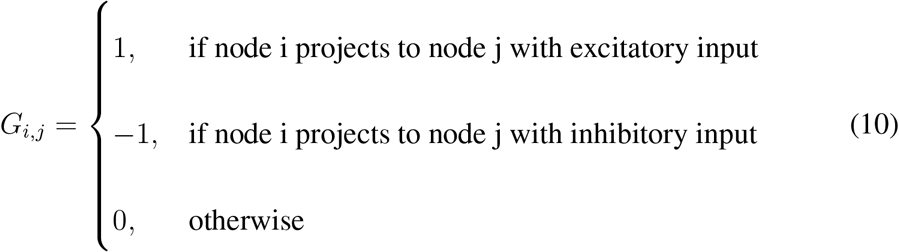

Then, we can use a 3-dimensional tensor to represent the false connections caused by common inputs. For example, if node *j* and node *k* receive common input from node *i*:

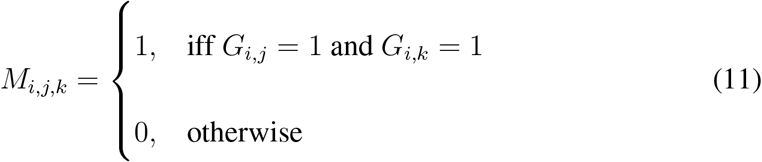

Therefore, we can compute a mask that labels all the false connections due to the confounder structure:

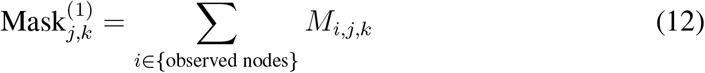

For the chain structure (e.g. node *i* projects to node *k*, then node *k* projects to node *j*), the mask is defined as:

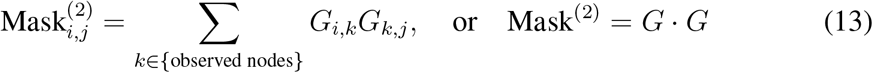

For the false connections caused by latent confounder structure, the mask is defined as:

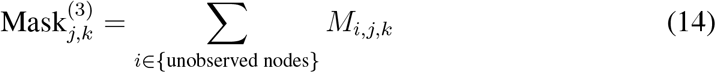

Lastly, Mask^(4)^ as the general false connection mask is defined as:

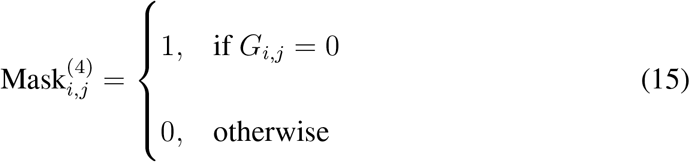

Given a certain mask Mask^(*k*)^ (*k* = 1, 2, 3, 4) and ground truth *G*, the true positive set (TP_*k*_) and the false positive set (FP_*k*_) is defined as:

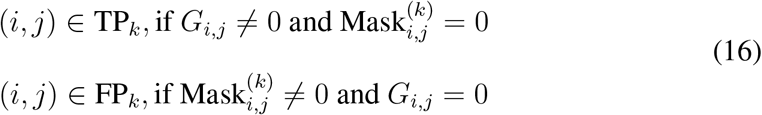

To quantify the performance of a functional connectivity estimation FC, we follow the convention of Smith et al. (2011) and use the “c-sensitivity” index defined in the paper. C-sensitivity is defined as the percentage of connections in TP that have a FC value which is above the 95% quantile of the FC values in FP:

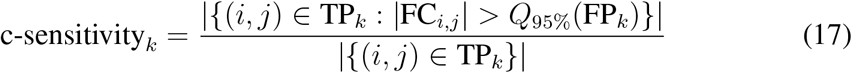

where (*i, j*) represents the connection from node *i* to node *j*, *|*FC_*i,j*_| is the absolute value of the functional connectivity from node *i* to node *j*, *Q*_95%_(FP_*k*_) represents the 95% quantile of the FC values in FP_*k*_ and *|{}|* is the number of elements in a given set.

## 3 Results

### 3.1 False connections in covariance-based methods

It is a well known problem that the commonly used covariance-based methods produce systematic false connections (Stevenson et al., 2008). Shown in Fig. 3A is the ground truth of the connection in our DCM model (Node No.1-50 are the observable nodes). Fig. 3B shows the results from the covariance method, Fig. 3C is the precision matrix, and Fig. 3D is the ICOV (sparsely regularized precision matrix). Comparing with the ground truth, all of these methods produce extra false connections. Here we briefly review the cause of these 3 types of false connections. The detailed mechanisms are well reviewed before (Stevenson et al., 2008).

*We define the diagonal strip of connections in the ground truth (Fig. 3A) as the +6 to +10 diagonal lines, because they are 6 to 10 steps away from the diagonal line of the matrix.*

Shown in Fig. 2A, confounder errors are produced because two nodes receive the same input from another node. The same input that passes into the two nodes generates positive correlation between the two nodes. However, there is no physiological connection between these two nodes. For example, as shown in Fig. 3A, node 1 projects to node 6 and node 7 (*A*_16_*, A*_17_ *>* 0), but node 6 and node 7 are not connected. However, because they share the same input from node 1, in Fig. 3B, there are positive connections on coordinate (6,7) and (7,6). Because this connection pattern is also applied to other nodes (node 2 projects to node 7 and node 8, node 3 projects to node 8 and so forth) there are false connections on the *±*1 to *±*4 diagonal lines of Fig. 3B.

Shown in Fig. 2B, the chain errors are due to the propagation of the covariance. Because node *A* connects to node *B* and node *B* connects to node *C*, the covariance method presents covariance between node *A* and node *C*, which do not have a physical connection. This phenomenon is shown in Fig. 3B as the extra diagonal strips.

Shown in Fig. 2C, the latent confounder errors are produced due to the common input passed into two nodes. However, in this case, they are from the unobserved latent inputs. For this DCM model, it is due to the inputs from the 10 latent nodes (node No.51-60) as shown on Fig. 3A. Because the latent nodes have broad connections to the observed nodes, they introduce a square box shape covariance pattern into Fig. 3B. And this type of false connections remains in the precision matrix and ICOV estimations (Fig. 3C, D) as the yellow color false connections on the bottom-left corner and the up-right corner.

### 3.2 Estimation from differential covariance method

Below we explain the estimation results from the differential covariance method. Comparing the ground truth connections in Fig. 4A with our final estimation in Fig. 4D, we see that our method essentially transformed the connections in the ground truth into a map of sources and sinks in a network. An excitatory connection, *i → j*, in our estimations have negative value for 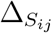 and positive value for 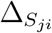, which means the current is coming out of the source *i*, and goes into the sink *j*. We note that there is another ambiguous case, an inhibitory connection *j → i*, which produces the same results in our estimations. Our method can not differentiate these two cases, instead, they indicate sources and sinks in a network.

**Figure 4:**
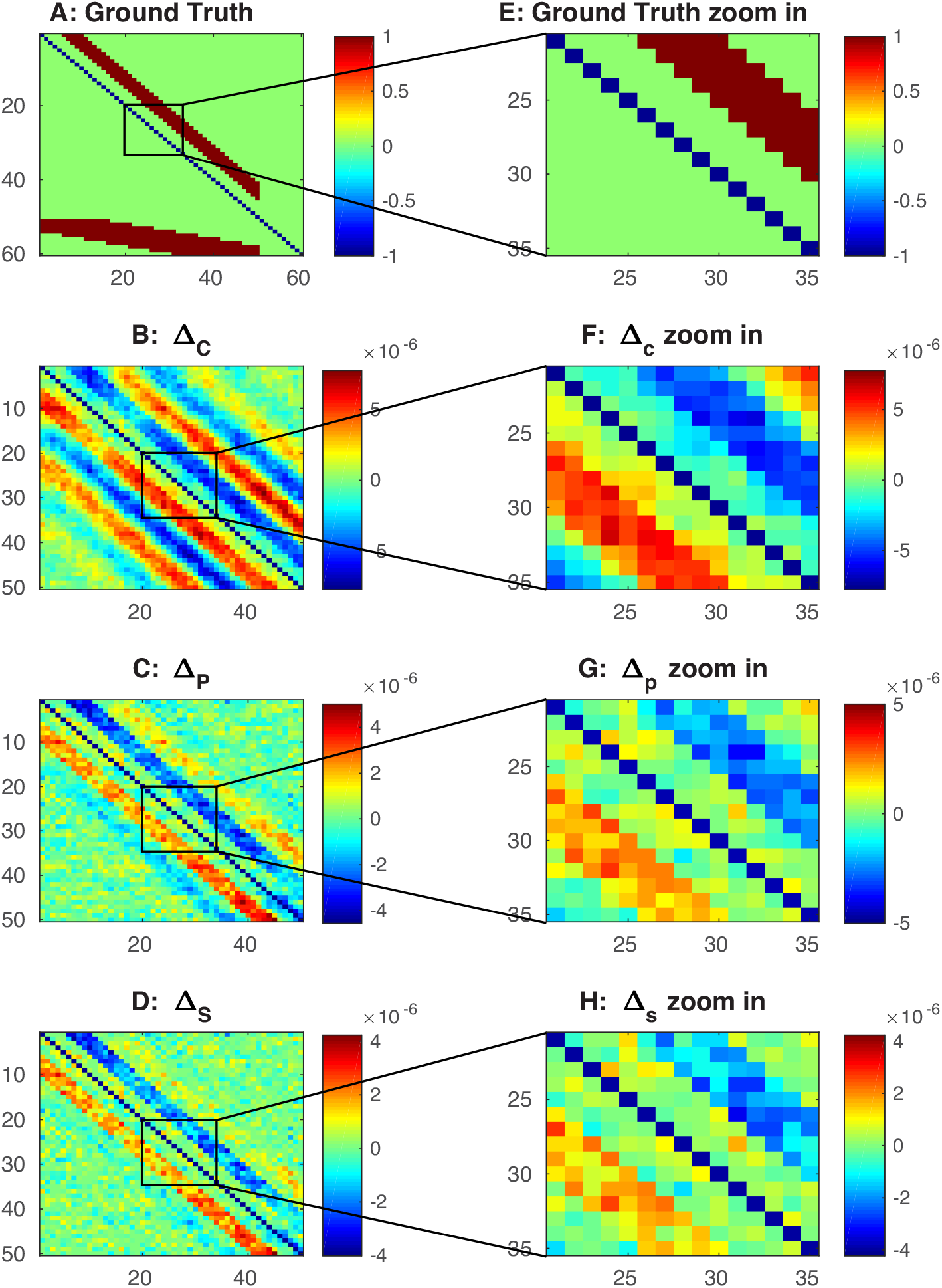
Differential covariance analysis of the DCM model. The color in B,C,D,F,G,H indicates direction of the connections. For element *A_i j_*, warm color indicates *i* is the sink, *j* is the source, i.e. *i ← j*, and cool color indicates *j* is the sink, *i* is the source, i.e. *i →j*. A) Ground truth connection matrix. B) Estimation from the differential covariance method (Δ_*C*_). C) Estimation from the partial differential covariance (Δ_*P*_). D) Estimation from the sparse+latent regularized partial differential covariance (Δ_*S*_). E) Zoom in of panel A. F) Zoom in of panel B. G) Zoom in of panel C. H) Zoom in of panel D.

#### 3.2.1 The differential covariance method reduces confounder errors

Comparing Fig. 3B and Fig. 4B, we see that confounder errors on *±*1 to *±*4 diagonal lines of Fig. 3B are reduced in Fig. 4B. This is quantified in Fig. 5A, where the differential covariance method has higher c-sensitivity score given confounder errors as the false positive set.

**Figure 5:**
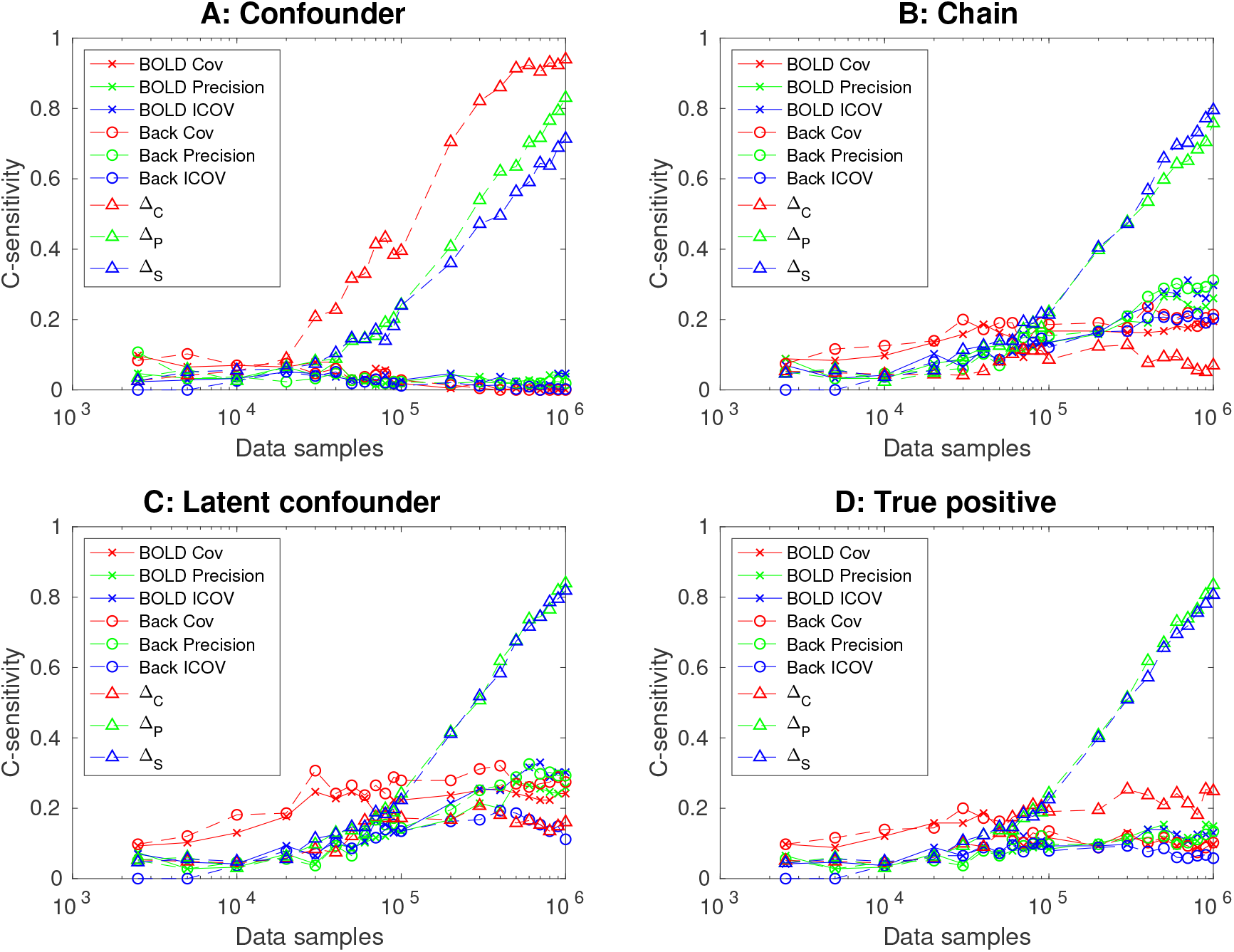
Performance quantification (c-sensitivity) of different methods with respect to their abilities to reduce the 3 types of false connections and their abilities to estimate the true positive connections using the dynamic causal modeling. The first three results (BOLD Cov, BOLD Precision and BOLD ICOV) are from covariance-based methods (covariance matrix, precision matrix and sparsely regularized precision matrix) directly applied to simulated BOLD signals. The next three results (Back Cov, Back Precision and Back ICOV) are from covariance-based methods applied to backward model reconstructed neural signals. The last three results (Δ_*C*_, Δ_*P*_ and Δ_*S*_) are from differential covariance methods applied to backward model reconstructed neural signals.

#### 3.2.2 The partial differential covariance reduces chain errors

Second, as we can see in Fig. 4B, there are propagation errors. By applying the partial covariance method, we regress out the interfering terms. Thus, there are fewer chain errors. Each method’s performance for reducing chain errors is quantified in Fig. 5B.

#### 3.2.3 The sparse+latent regularization reduces latent confounder errors

Third, we use the sparse latent regularization to remove the covariance introduced by the latent variables. As mentioned in the method section, when the observable nodes’ connections are sparse and the number of latent inputs is small, covariance introduced by the latent inputs can be separated. As shown in Fig. 4D, the external covariance in the background of Fig. 4C is reduced, while the true positive connections and the directionality of the connection is maintained. Each method’s ability to reduce this type of error is quantified in Fig. 5C, and each method’s overall performance at reducing all errors is quantified in Fig. 5D.

### 3.3 Thalamocortical model’s results

In addition to the DCM simulations, we also benchmarked various methods with a more realistic Hodgkin-Huxley based model.

The covariance-based methods are applied to both the raw BOLD signals and the reconstructed neural signals. The differential covariance based methods are applied to both the raw BOLD signals and the reconstructed neural signals as well. Compared to the ground truth in Fig. 6A, covariance-based estimations failed to show the true connections. Also confounder errors are noticeable on the *±*1 diagonal line of Fig. 6 C and D. We show later that, applying the covariance-based methods to the reconstructed neural signals does provide better results, however, covariance-based methods still suffer from the 3 types of false connections mentioned above.

On the other hand, by applying differential covariance to the reconstructed neural signals, we see in Fig. 7B, Δ_*C*_ provides directionality information of the connection. However, Δ_*C*_ estimation is contaminated by connections introduced from the propagation of the signals and the latent inputs. Confounder errors are greatly reduced. In Fig. 7C, the partial covariance reduces chain errors and produces Δ_*P*_. Finally, in Fig. 7D, the sparse+latent regularization is applied to reduce latent confounder errors.

**Figure 7:**
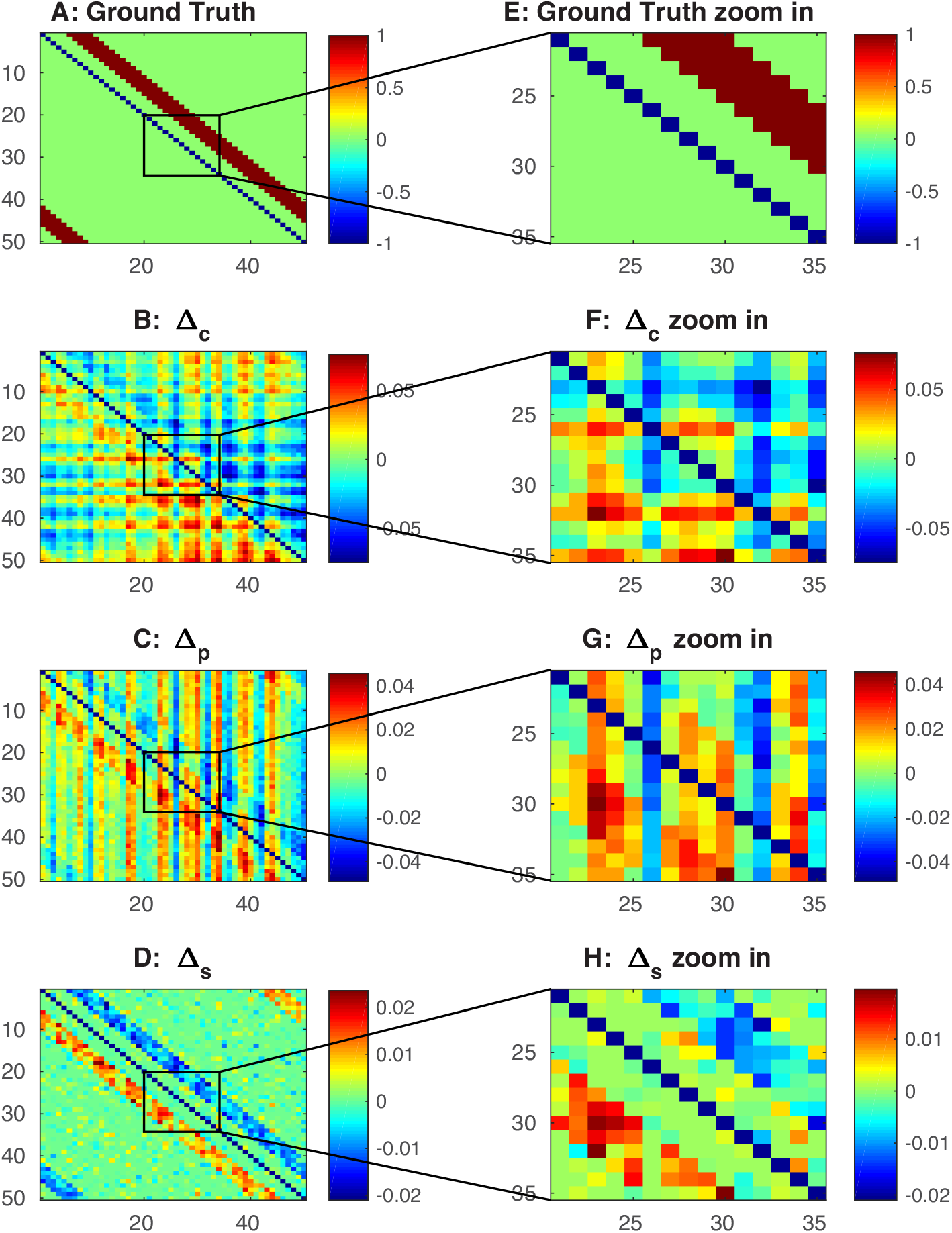
Differential covariance analysis of the thalamocortical model’s reconstructed neural signals. The color in B,C,D,F,G,H indicates direction of the connections. For element *A_ij_*, warm color indicates *i* is the sink, *j* is the source, i.e. *i ← j*, and cool color indicates *j* is the sink, *i* is the source, i.e. *i → j*. A) Ground truth connection matrix. B) Estimation from the differential covariance method. C) Estimation from the partial differential covariance method. D) Estimation from the sparse+latent regularized partial differential covariance method. E) Zoom in of panel A. F) Zoom in of panel B. G) Zoom in of panel C. H) Zoom in of panel D.

In Fig. 8, each estimator’s performance in reducing different types of false positive connections is quantified. As explained above, while covariance-based methods perform well at chain errors and latent confounder errors, differential covariance method performs better at reducing confounder errors. Thus it achieves better performance overall.

**Figure 8:**
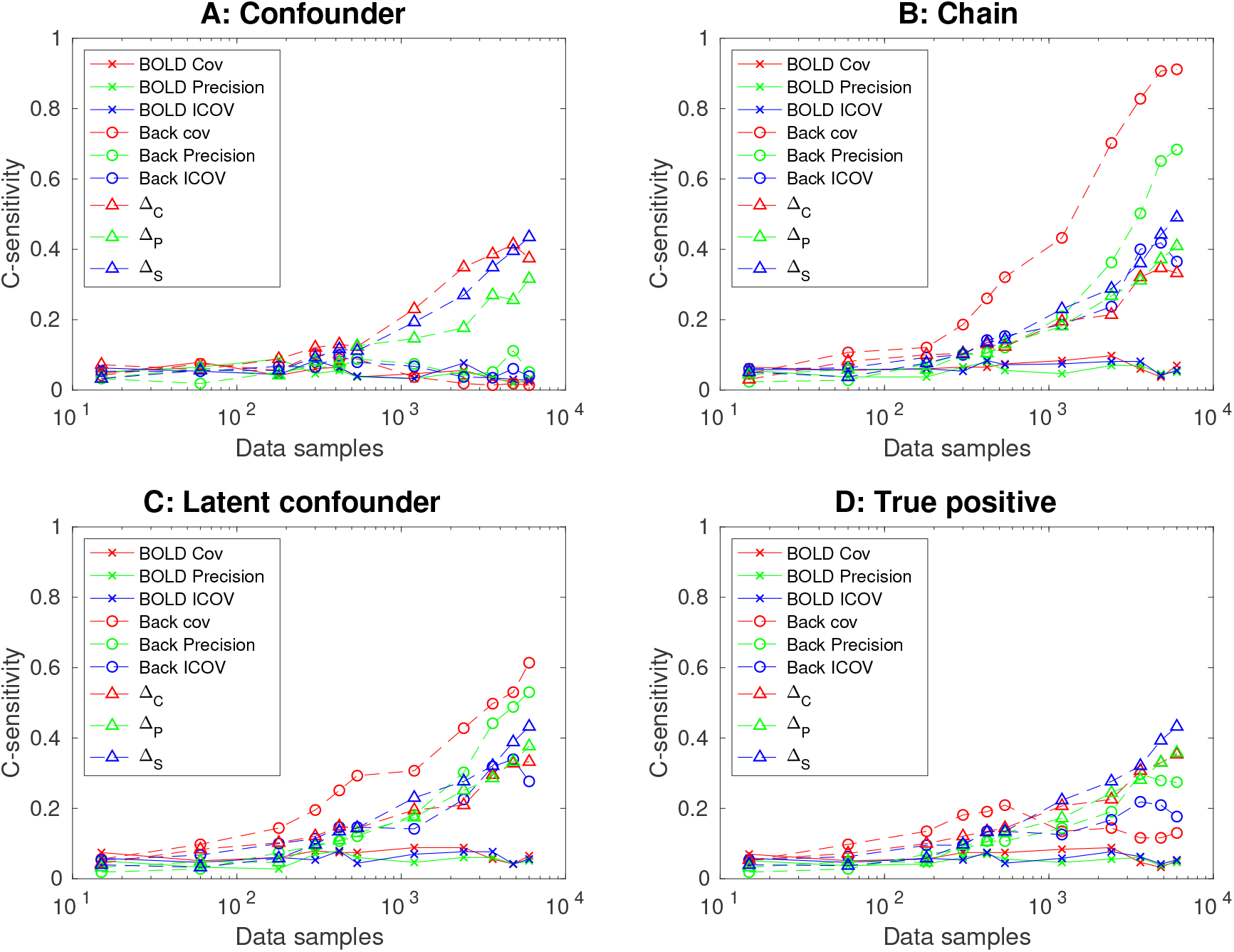
Performance quantification (c-sensitivity) of different methods with respect to their abilities to reduce the 3 types of false connections and their abilities to estimate the true positive connections using the thalamocortical model. The first three results are from covariance-based methods directly applied to simulated BOLD signals. The next three results are from covariance-based methods applied to backward model reconstructed neural signals. The last three results are from differential covariance methods applied to backward model reconstructed neural signals.

### 3.4 The backward model and the differential covariance are both needed to produce correct estimations

Above, we explained our newly developed backward model and the differential covariance method. Below we show that both steps are necessary to produce good estimation. The purpose of the backward model is to restore the dynamic form of neural signals, such that it is suitable for the differential covariance estimation. As shown in Fig. 9, if we directly apply the differential covariance method to the raw BOLD signals, the estimator’s performance is poor. The performance quantified using c-sensitivity was shown in Fig. 8. The backward model is a better procedure (Appendix. E) to reconstruct neural signal compared to blind deconvolution (Wu et al., 2013), which is commonly used to deconvolute resting state fMRI signals. In addition, the backward model is robust to parameter jittering because the same set of backward parameters could still reconstruct the signals simulated using forward Balloon model with a wide range of parameters (Appendix. F).

**Figure 9:**
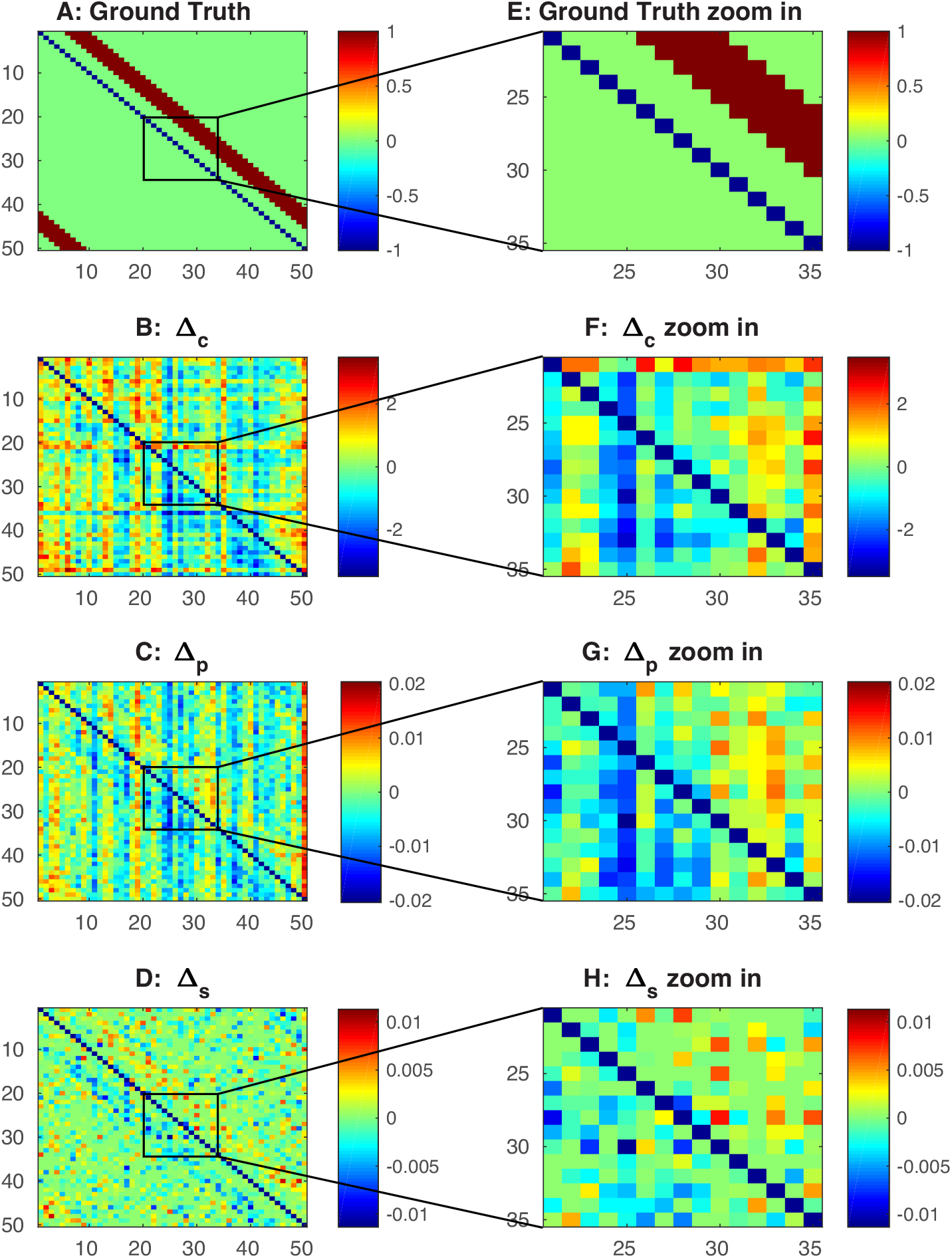
Differential covariance method applied directly to simulated BOLD signals without applying the backward model. The color in B,C,D,F,G,H indicates direction of the connections. For element *A_ij_*, warm color indicates *i* is the sink, *j* is the source, i.e. *i ← j*, and cool color indicates *j* is the sink, *i* is the source, i.e. *i → j*. A) Ground truth connection matrix. B) Estimation from the differential covariance method. C) Estimation from the partial differential covariance method. D) Estimation from the sparse+latent regularized partial differential covariance method. E) Zoom in of panel A. F) Zoom in of panel B. G) Zoom in of panel C. H) Zoom in of panel D.

On the other hand, the differential covariance method is also needed to reduce the false connections. In Fig. 10, estimations from the covariance-based methods applied to the reconstructed neural signals failed to reduce the false connections mentioned above. Particularly, confounder errors that are on the *±*1 and *±*2 diagonal lines are apparent in Fig. 10 C and D. This comparison can also be seen from the quantifications on Fig. 5 and Fig. 8. For both simulations, even using the same backward processed signals, the differential covariance method performs better at reducing confounder errors than the covariance-based methods. While the covariance-based methods perform well at reducing chain errors and latent confounder errors, differential covariance method’s performance is better overall. Therefore, the differential covariance method is needed to make better estimations after the backward reconstruction step.

**Figure 10:**
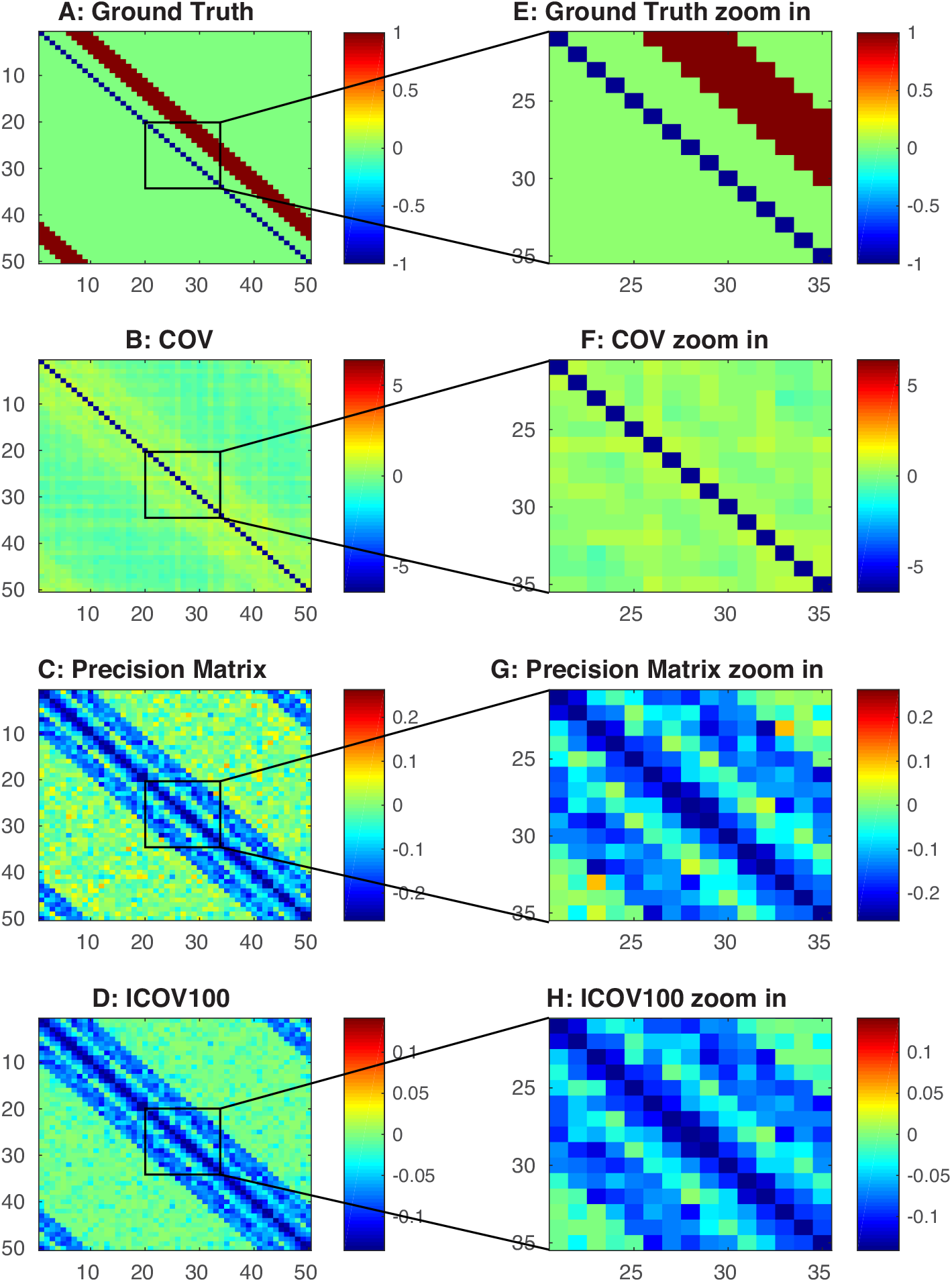
covariance-based methods applied to backward model reconstructed signals. A) Ground truth connection matrix. B) Estimation from the covariance method. C) Estimation from the precision matrix method. D) Estimation from the sparsely regularized precision matrix (ICOV) method. E) Zoom in of panel C. F) Zoom in of panel D. E) Zoom in of panel A. F) Zoom in of panel B. G) Zoom in of panel C. H) Zoom in of panel D.

### 3.5 Benchmark using previous study’s dataset

To show that our new method is generalizable outside of the two example simulations just given, we benchmarked various methods using a previously generated dataset containing 28 BOLD signal simulations (Smith et al., 2011). Top performers mentioned in the previous study (Smith et al., 2011) (ICOV, Precision matrix, Bayes net, and Patel’s *κ*) are included as benchmarks. Shown in Fig. 11A, For the thalamocortical ionic model, we see that the Δ_*S*_ outperforms others. This is mainly because the method’s better performance at reducing confounder errors.

**Figure 11:**
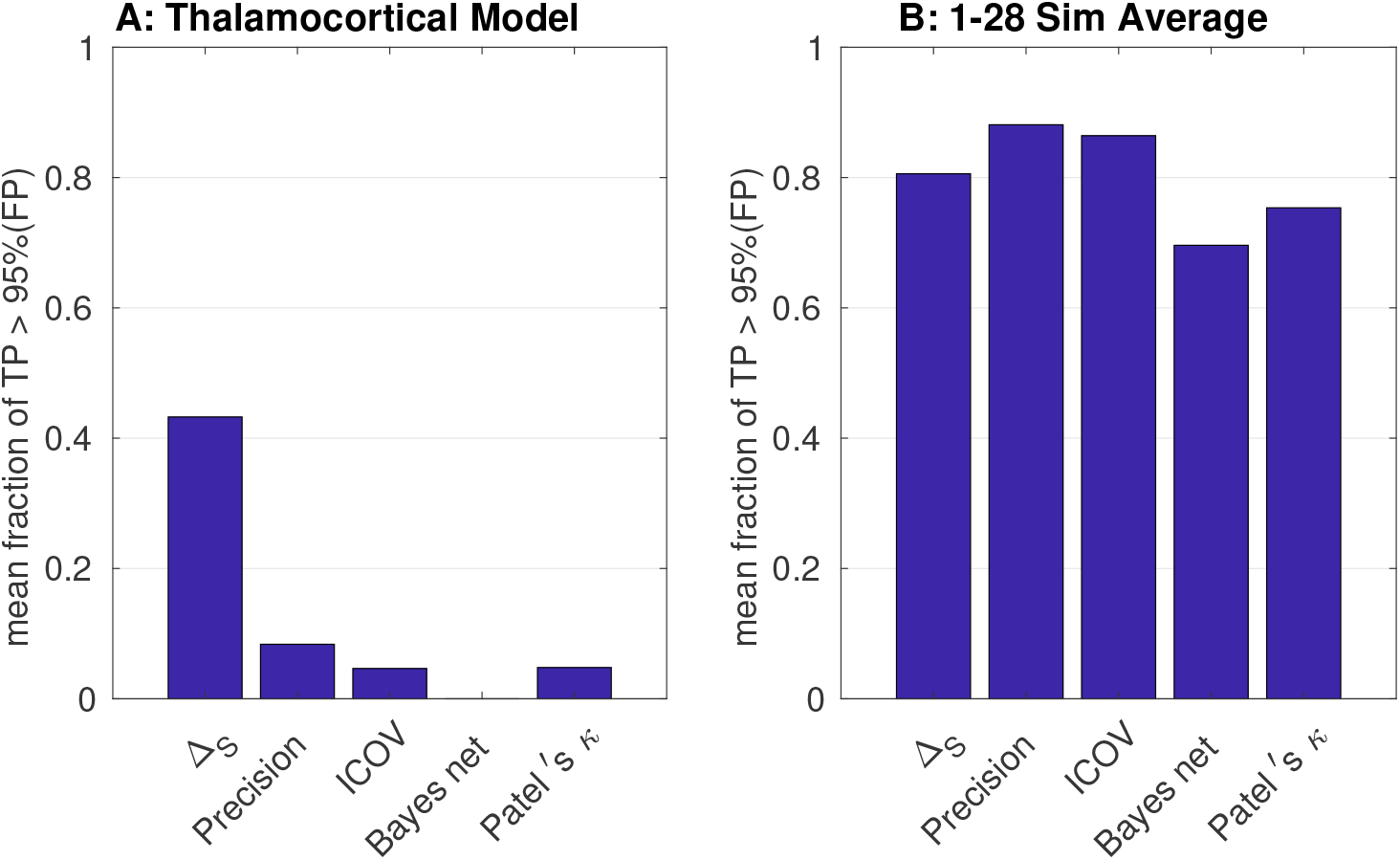
A) Performances (c-sensitivity) of different methods on the thalamocortical model simulations. B) Average performances across the 28 datasets from a previous study(Smith et al., 2011). *The Bayes net column refers to the average performance of all Bayes net methods mentioned in Appendix A.2.* Method’s performance on individual data sets is provided in Appendix A.5.

For the 28 simulations from the previous study (Smith et al., 2011), the differential covariance method’s average performance is comparable to other top methods identified previously (Fig. 11B). Performance on individual data sets is provided in Appendix A.5. Because most of these 28 simulations only have 5 nodes and the connection matrix is not sparse, we dropped the sparse+latent regularization step in our method, except for simulation No.4, the only one that has a sparse connection matrix (50 nodes, about 70 connections).

## 4 Discussion

### 4.1 Main results

*The goal of this paper was to reconstruct neural signals from BOLD signals. We used synthetic BOLD data from detailed neural models to benchmark a novel differential covariance method to estimate the underlying functional connectivity. Although the relationship between fMRI and neuronal activity is a well-studied topic (for reviews, see Heeger and Ress (2002); Logothetis (2002)), we found that a custom-designed backward model was needed for our differential covariance method to to efficiently and validly recover the underlying ground truth (Fig. 9). This new method not only reduced estimation errors made by previous methods, but also provided information about the directionality of the connections.*

For the DCM dataset from a previous study (Smith et al., 2011), differential covariance had similar performance to the previously benchmarked best methods (covariance-based methods). As explained above, each step of the differential covariance method is specialized at reducing one type of false connections. However, since the presence of these 3 types of false connections are minimal in this dataset, our new method does not outperform the covariance-based methods. Further, because the size of the network is small, the connectivity matrix is not sparse, so we cannot take advantage of the sparse+latent regularization step in our method. We believe this will not be an issue when applying differential covariance method to real-world problems, because the brain network has a much higher dimensionality and is sparser.

### 4.2 Caveats and future directions

As mentioned above, we made several key assumptions that should be further tested and tuned in the future. Our approach depends on the neural signals reconstructed from the fMRI transfer function. Even though the fMRI transfer function we used (the Balloon model) is based on the fundamental vesicular dynamics, whether high quality neural signals can be reconstructed from the noisy real fMRI instruments should be further studied. *The backward reconstruction process also had lower noise tolerance (Appendix. D). In addition, the sparse and low-rank assumptions were critical to the cases we simulated. These assumptions should be validated in applications to fMRI recordings.*

The next step is to evaluate the performance of differential covariance analysis on fMRI recordings and compare the results with other methods for measuring functional connectivity. Preliminary results from analyzing fMRI resting state data from humans and mice with this new method are promising and will be reported elsewhere. Estimating functional connectivity is of great importance in understanding the dynamical patterns of brain activity and differential covariance analysis of fMRI recordings offers an new way to identify causal interactions between brain region.

## Acknowledgement

This research was supported by the Office of Naval Research (N00014-16-1-2829) and NIH/NIBIB (R01EB026899).

## Appendix

## A Previous methods

In this section we review several top performing methods benchmarked in a previous study (Smith et al., 2011).

## A.1 covariance-based methods

The well used covariance-based methods are used as a benchmark in this paper because a prior work (Smith et al., 2011) has shown its superior performance by applying to simulated BOLD signal.

## A.1.1 covariance method

The covariance matrix is defined as:

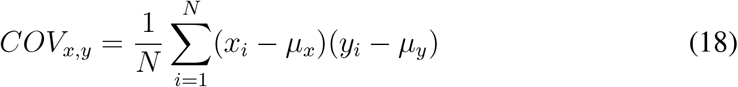

Where *x* and *y* are two variables. *μ*_*x*_ and *μ*_*y*_ are their population mean.

## A.1.2 Precision Matrix

The precision matrix is the inverse of the covariance matrix:

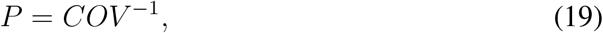

It can be considered as one kind of partial correlation. Here we briefly review this derivation, because we use it to develop our new method. The derivation here is based on and adapted from Cox and Wermuth (1996).

We begin by considering a pair of variables (*x, y*), and remove the correlation in them introduced from a control variable *z.*

First, we define the covariance matrix as:

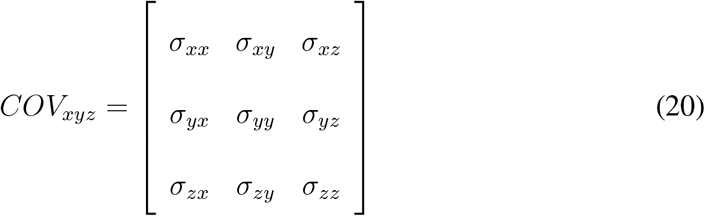

By solving the linear regression problem:

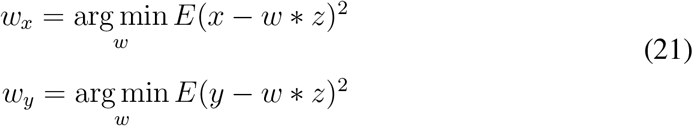

we have:

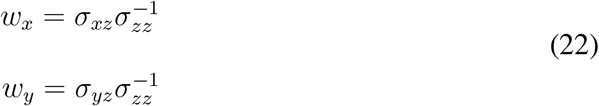

then, we define the residual of *x, y* as,

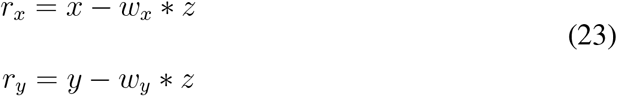

Therefore, the covariance of *r*_*x*_, *r*_*y*_ is:

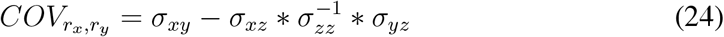

On the other hand, if we define the precision matrix as:

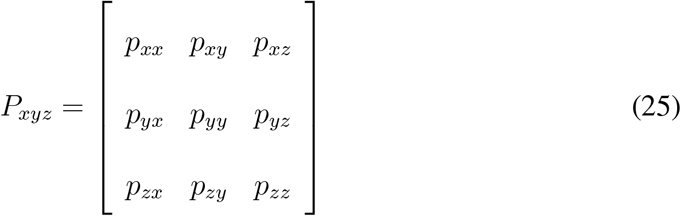

Using Cramer’s rule, we have:

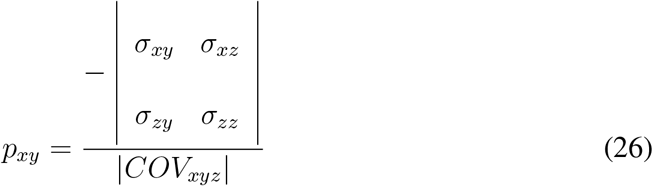

Therefore,

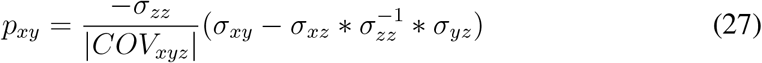

So *p*_*xy*_ and 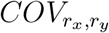 are differed by a ratio of 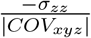.

## A.1.3 Sparsely regularized precision matrix

If the precision matrix is expected to be sparse, one way to efficiently and accurately estimate this matrix, especially when the dimension is large, is regularize the estimation using the Lasso method (Hsieh et al., 2014, 2013; Friedman et al., 2008; Banerjee et al., 2006).

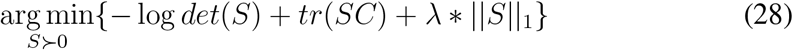

where C is the sample covariance matrix.

We used the implementation (https://www.cs.ubc.ca/~schmidtm/Software/L1precision.html) mentioned in the previous work (Smith et al., 2011) to benchmark our new method.

## A.2 Bayes-net method

A variety of the Bayes Net estimation methods are implemented in the Tetrad toolbox (http://www.phil.cmu.edu/tetrad/). Following the convention of the prior work (Smith et al., 2011), we tested CCD, CPC, FCI, PC and GES. PC (”Peter and Clark”) is a causal inference algorithm used to estimate the causal relationship in a directed acyclic graph (DAG) (Meek, 1995). CPC (Conservative PC) was based on the PC algorithm, but produce less false connections (Ramsey et al., 2012). GES (Greedy Equivalence Search) is another implementation based on the PC algorithm, but it uses a score system to estimate the causal connections (Chickering, 2003; Ramsey et al., 2010). The FCI (Fast Causal Inference) algorithm is a different approach, which takes into account the presence of latent confounder and selection bias (Zhang, 2008). CCD (Cyclic Causal Discovery) unlike other algorithms mentioned, allows cycles in the network for estimation (Richardson and Spirtes, 1996). The code used in this paper is based on the Tetrad Toolbox and further adapted by the authors of Smith et al. (2011) for fMRI data.

## A.3 Patel’s κ method

The Patel’s *κ* measurement assess the relationship between pairs of distinct brain regions by comparing expected joint and marginal probabilities of elevated activity of voxel pairs through a Bayesian paradigm (Patel et al., 2006). For this paper, we used the implementation generously provided by the authors of Smith et al. (2011);

## A.4 Sparse latent regularization

Prior studies(Banerjee et al., 2006; Friedman et al., 2008) have shown that regularizations can provide better estimation if the ground truth connection matrix has a known structure (e.g. sparse). For all data tested in this paper, the sparse latent regularization(Yatsenko et al., 2015) worked best. For a fair comparison, we applied the sparse latent regularization to both the precision matrix method and our differential covariance method.

In the original sparse latent regularization method, people made the assumption that a larger precision matrix *S* is the joint distribution of the *p* observed neurons and *d* latent neurons (Yatsenko et al., 2015). i.e.

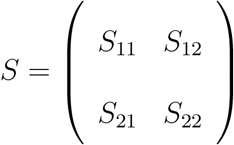

Where *S*_11_ corresponds to the observable neurons. If we can only measure the observable neurons, the partial correlation computed from the observed neural signals is,

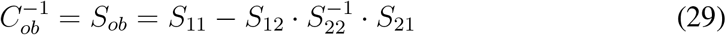

because the latent latent neurons as shown in Eq. 29 introduce correlations into the measurable system. We denote this correlation introduced from the latent inputs as

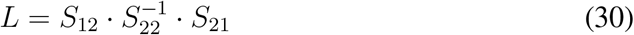

If we can make the assumption that the connection between the visible neurons are sparse, i.e. *S*_11_ is sparse and the number of latent neurons is much smaller than the number of visible neurons, i.e. *d* << p. Then, prior works (Chandrasekaran et al., 2011) have shown that if *S*_ob_ is known, *S*_11_ is sparse enough and L’s rank is low enough (within the bound defined in Chandrasekaran et al. (2011)), then the solution of

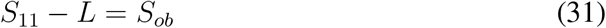

is uniquely defined and can be solved by the following convex optimization problem

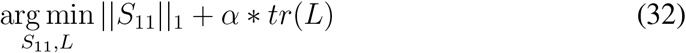

under the constraint that

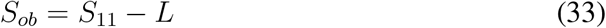

Here, *|| ||*_1_ is the L1-norm of a matrix, and *tr*() is the trace of a matrix. *α* is the penalty ratio between the L1-norm of *S*_11_ and the trace of L and is set to 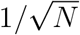 for all our estimations.

However, the above method is used to regularize precision matrix. For our differential covariance estimation, we need to make small changes to the derivation. Note that if we assume the neural signals of the latent neurons are known, and let *l* be the indexes of these latent neurons, then from our previous section (section 2.2.3),

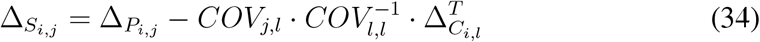

removes the *V*_*latent*_ terms.

Even if *l* is unknown,

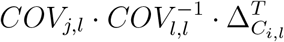

is low-rank, because it is bounded by the dimensionality of *COV*_l,l_, which is *d*. And Δ_*S*_ is the internal connections between the visible neurons, which should be a sparse matrix. Therefore, letting

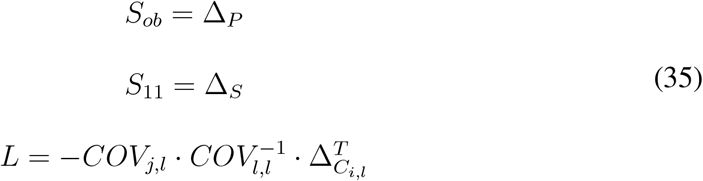

we can use the original sparse+latent method to solve for Δ_*S*_. In this paper, we used the inexact robust PCA algorithm (http://perception.csl.illinois.edu/matrix-rank/sample_code.html) to solve this problem (Lin et al., 2011).

## A.5 Methods’ performance for individual datasets

In Fig.12, the performance of different methods mentioned above is quantified using c-sensitivity on individual datasets simulated in Smith et al (Smith et al., 2011). For comparison, the methods’ performance on thalamocortical model and their average performance across all 28 data sets are shown in the first two columns.

**Figure 12:**
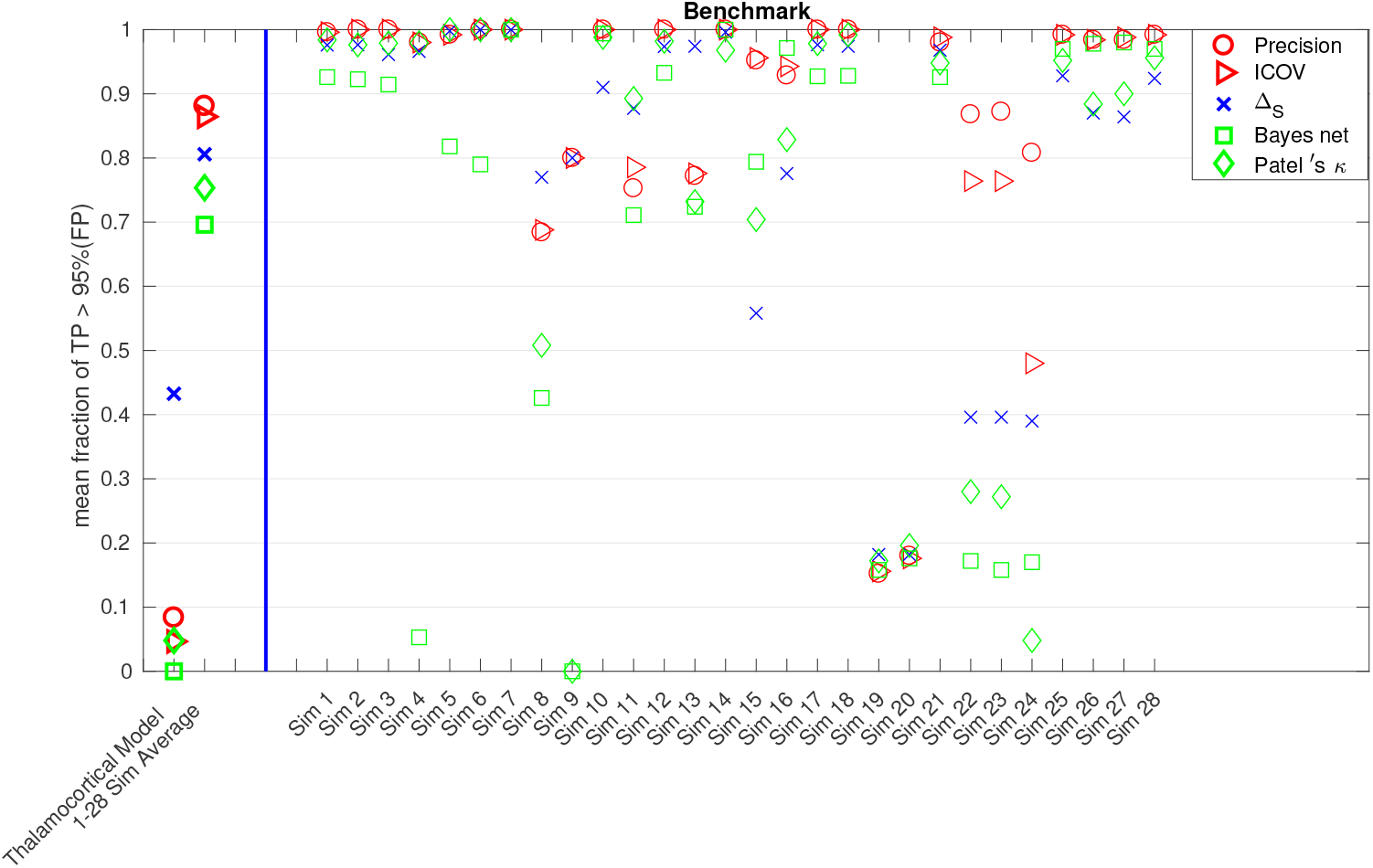
The first column is method’s performance (c-sensitivity) on our more realistic thalamocortical ionic model.The second column is method’s average performance across the 28 data sets from a previous study(Smith et al., 2011). Column 3-30 are method’s performances for each of the 28 data sets.

## B Balloon Model Parameters

The values and notations for parameters involved in the forward Balloon model (Eq.2) are shown in Table.1. The values of the coefficients derived from these parameters are evaluated in Eq.36.

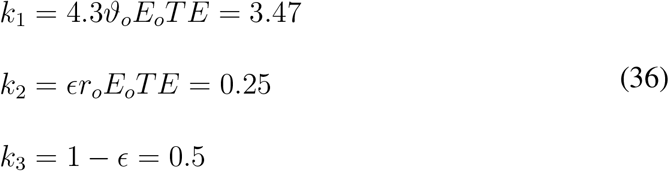

**Table 2:**
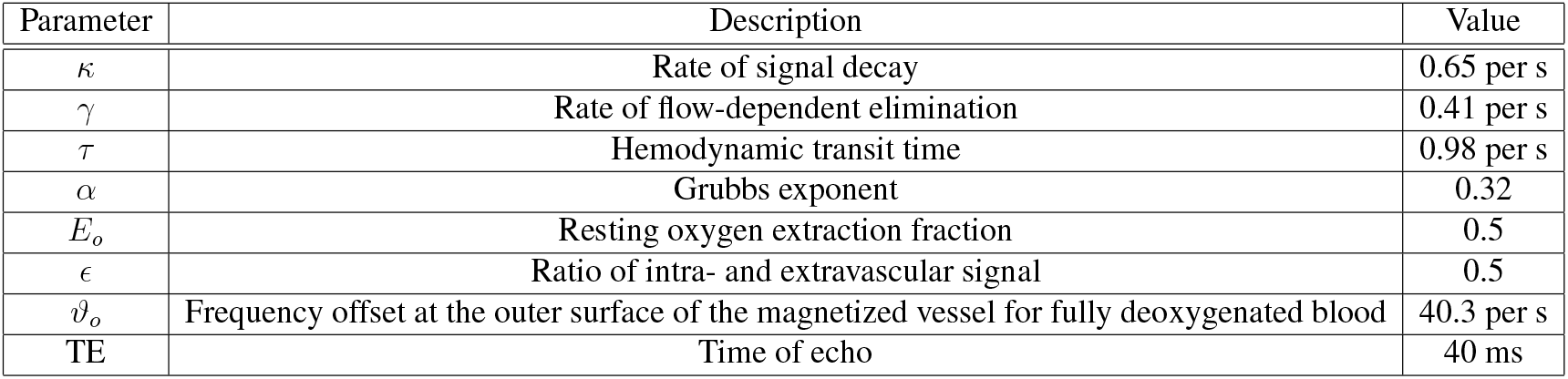
Backward reconstruction parameters

## C Backward Model Parameters

The values and notations for parameters involved in the backward reconstruction model (Eq.3) are shown in Table.2. The values of the coefficients (Eq.4) derived from these parameters are evaluated in Eq.37.

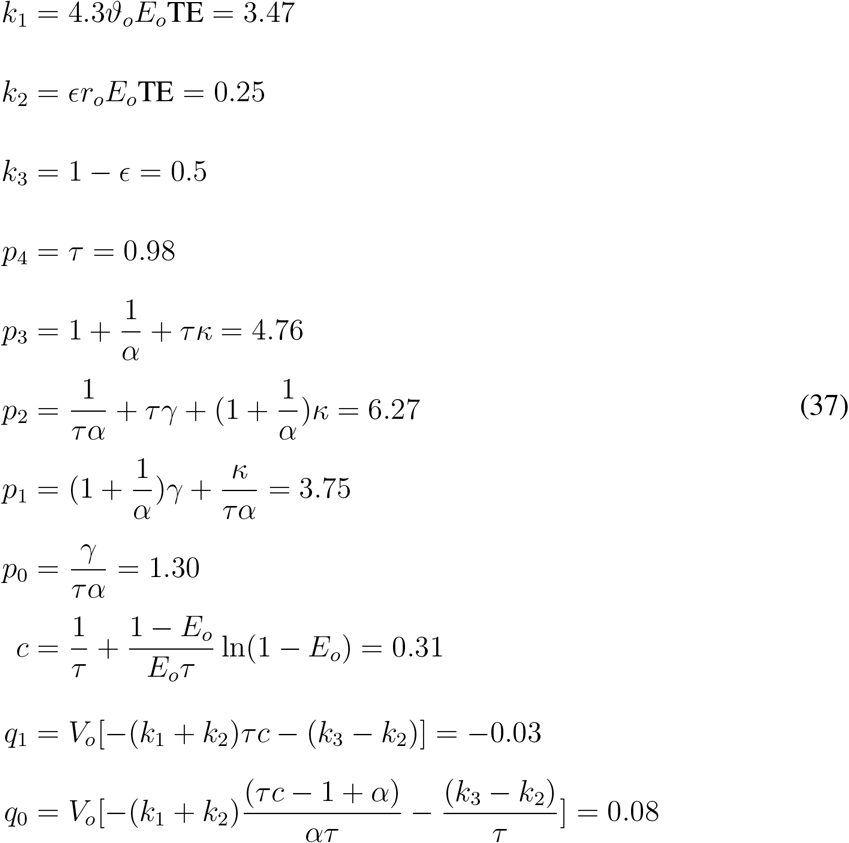

## D Noise Tolerance

When using numerical differentiation method to compute the reconstructed signal in Eq.4, we noticed that noise tolerance of the method depends on the sampling frequency. For example, if white noise 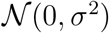 was introduced into BOLD signal *y* sampled at 10Hz, the noise was magnified by 10 times for 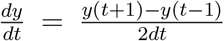 because noise variance in the nominator is 2*σ*^2^ and *dt* = 0.1. The observation was confirmed in Fig.13 where differential covariance analysis was applied to both noisy signal (right panel) and clean signal (left panel) after backward reconstruction. While we could still identify the overall connection structure from noisy signal sampled at 1Hz, the connection pattern was totally destroyed for noisy signal sampled at 10Hz.

**Figure 13:**
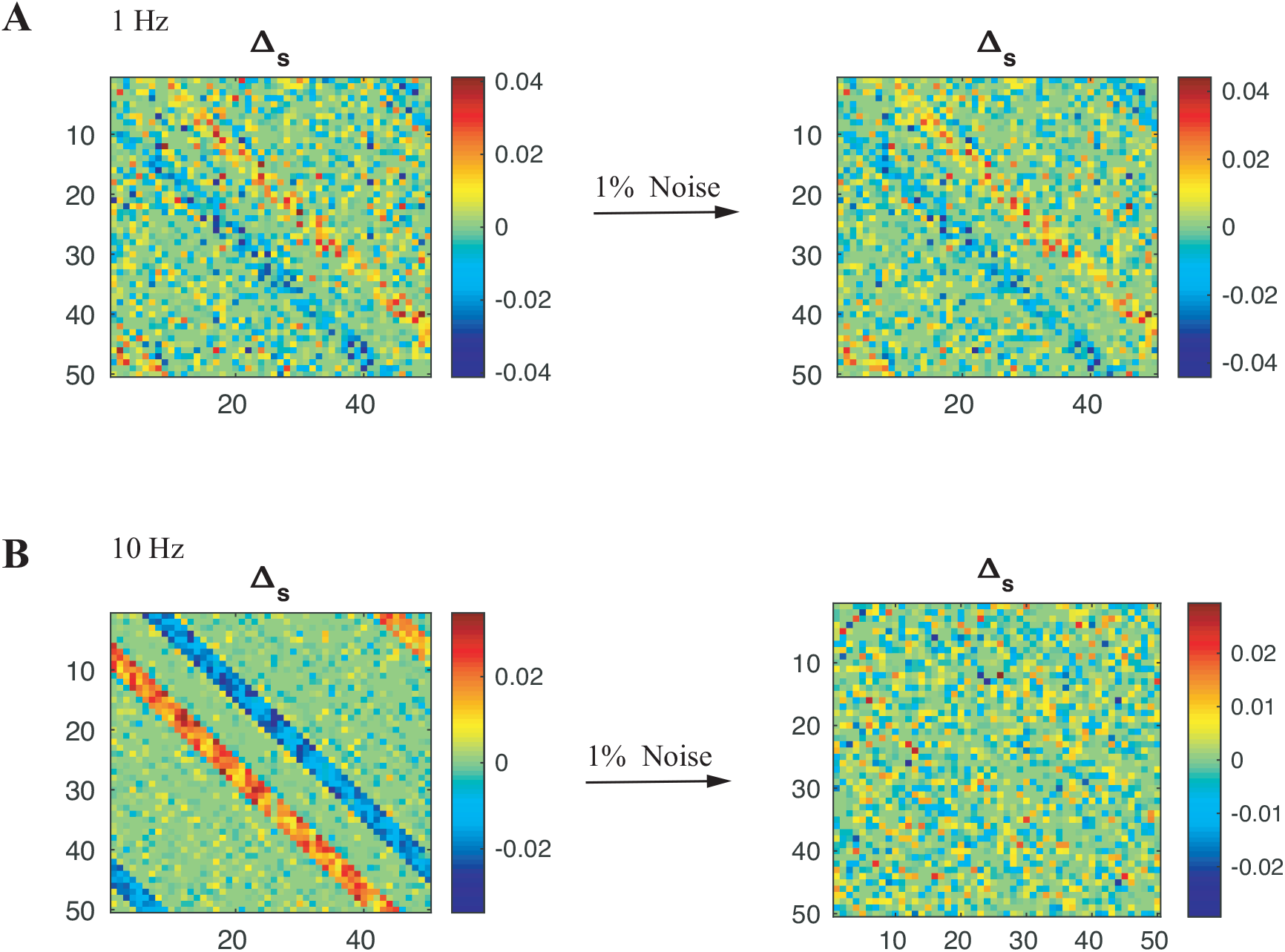
Noise tolerance of the method. BOLD signals were simulated from the thalamacortical model. A) left panel: Δ_*S*_ estimated from the clean signal sampled at 1Hz; right panel: Δ_*S*_ estimated from the noisy (white noise with variance equals to 1% of the signal level) signal sampled as 1Hz. B) left panel: Δ_*S*_ estimated from clean signal sampled as 10Hz; right panel: Δ_*S*_ estimated from noisy (1%) signal sampled at 10Hz

## E Benchmark with blind deconvolution

Blind deconvolution aims to reduce the confounding effects of hemodynamic response function (HRF) on resting state fMRI signals. For task-related fMRI, neural population dynamics can be captured by modeling signal dynamics with explicit exogenous inputs, which is absent during resting state fMRI recordings. Blind deconvolution considers resting state fMRI as ‘spontaneous event related’ and then estimates region-specific HRF and performs deconvolution on the signal. More details could be found in (Wu et al., 2013).

Here we performed blind deconvolution on BOLD signals simulated from thalamocortical model and applied differential covariance analysis on the deconvoluted signal (Fig. 14). The performance of blind deconvolution is not as good as the backward reconstruction process.

**Figure 14:**
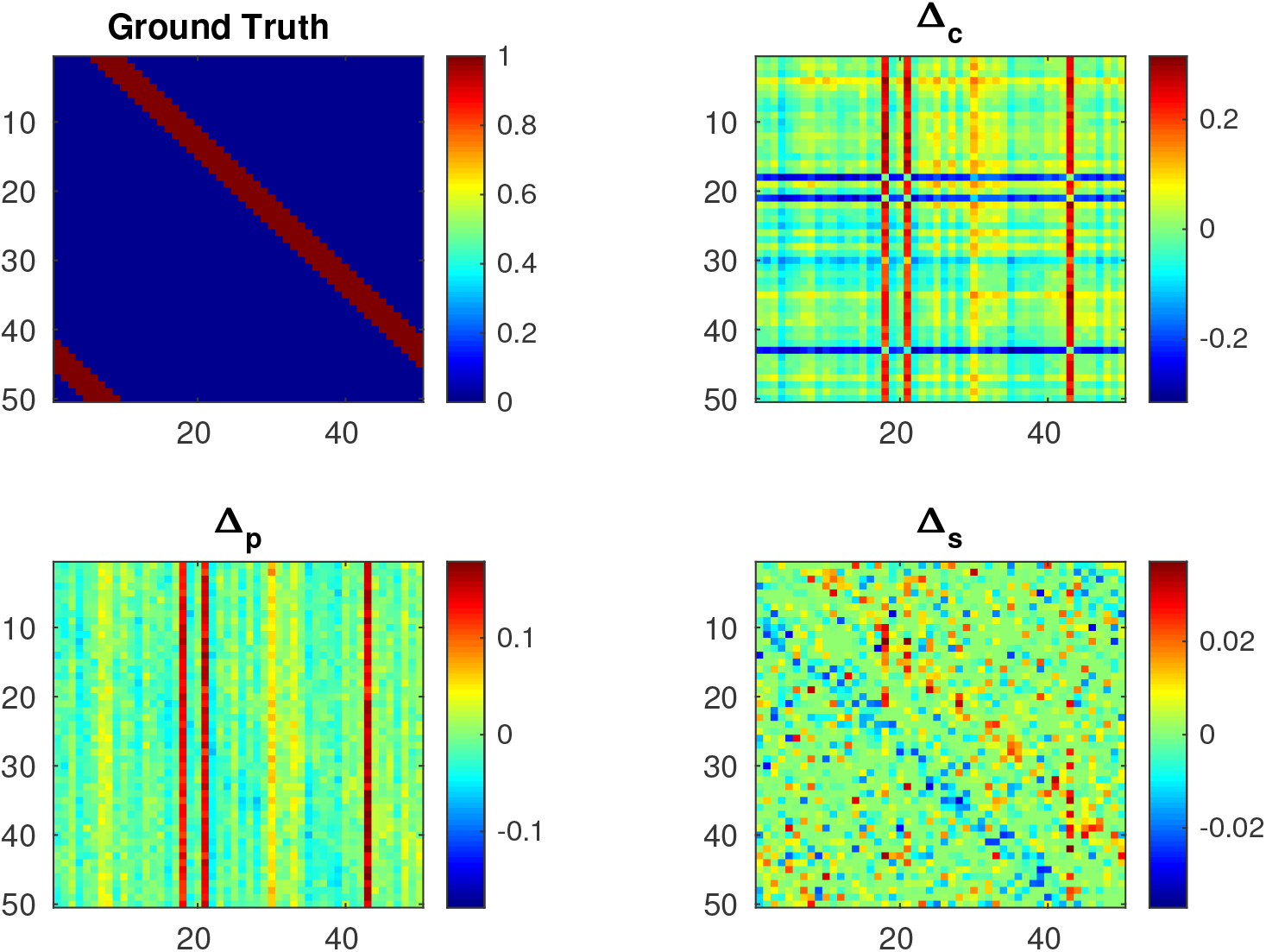
Differential covariance analysis applied to blind deconvoluted BOLD signal simulated from the thalamocortical model.

## F Robustness of the backward model

There is considerable uncertainty regarding *E*_o_ and *ϵ* (Stephan et al., 2007). To evaluate whether the backward model is applicable to different parameter values of *E*_o_ and *ϵ*, we randomly sampled *E*_o_ and *ϵ* from the following distribution (Eq. 38) and simulated multiple sets of BOLD signals. Backward reconstruction with *E*_o_ = 0.5 and *ϵ* = 0.5 followed by differential covariance analysis was applied to the signals and the performance of Δ_*s*_ was quantified by c-sensitivity (Fig. 15). The c-sensitivity of Δ_*S*_ remains to be high with respect to a wide range of forward Balloon model parameters.

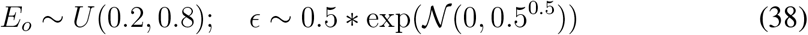

**Figure 15:**
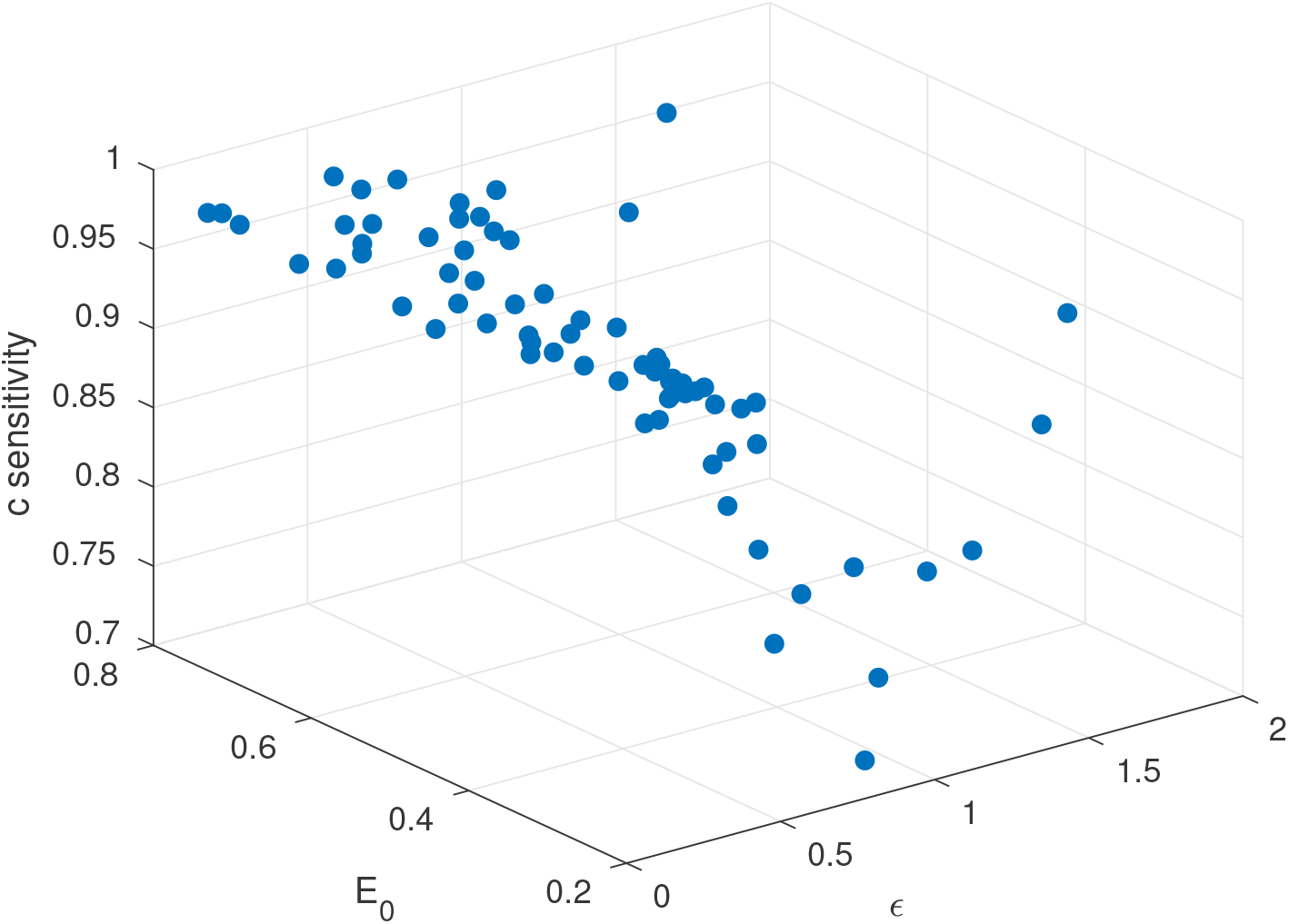
C-sensitivity of Δ_*S*_ with different forward Balloon model parameters

